# Tracking emotional interference over time: Differential effects of dorsolateral and ventromedial prefrontal stimulation

**DOI:** 10.64898/2026.08.02.742381

**Authors:** T. Feutren, V. Braud, L. Fabre

## Abstract

Although a substantial body of evidence has demonstrated that emotional interference affects cognitive performance, relatively little is known about its temporal evolution and the respective brain regions underlying its regulation. The present study investigated the temporal dynamics of emotional interference and the contribution of prefrontal regions involved in its regulation. Forty-eight participants completed a 2-back task and a Set-switching task under neutral and negative emotional conditions while receiving sham, dorsolateral prefrontal cortex (dlPFC), or ventromedial prefrontal cortex (vmPFC) stimulation. Stimulation was administered either online during task performance or after a 5 min pre-task period. Consistent with previous findings, negative emotions impaired executive performance, particularly during high-demand updating conditions. Critically, time-resolved analyses revealed that emotional interference evolved dynamically throughout task performance and was differentially modulated by prefrontal stimulation. The most consistent stimulation effects emerged after approximately 10 minutes of cumulative stimulation exposure and varied as a function of the stimulation site, executive-control demands, and stimulation timing. Notably, online stimulation produced more consistent modulation than pre-task stimulation. Together, these findings indicate that both emotional interference and its neuromodulation are dynamic processes. More broadly, they suggest that the contribution of prefrontal control systems to emotion-cognition interactions may be better understood through their temporal evolution rather than through static measures of performance alone.

## Introduction

Emotional interference refers to the disruptive influence of emotional stimuli on ongoing cognitive processing. Previous studies have consistently shown that emotional interference impairs performance across a wide range of cognitive functions, including attention, working memory, and executive control, as emotionally salient stimuli tend to receive prioritized processing and compete with ongoing goal-directed behaviors [1,2,3,4]. Although the behavioral consequences of emotional interference are well documented, the cognitive mechanisms through which emotional information influences executive control processes remain poorly understood. In particular, little is known about how emotional interference is regulated over time and whether this regulation relies on dynamic changes in cognitive control mechanisms.

Previous findings further suggest that emotional interference is not a static phenomenon. Repeated exposure to emotional distractors has been associated with progressive reductions in interference, reflecting habituation and adaptive attentional filtering mechanisms [5,6]. Moreover, the magnitude of emotional interference depends on contextual factors such as cognitive load and distractor frequency [7,8]. Together, these findings suggest that emotional interference relies on dynamic processes that evolve throughout task performance.

However, previous studies have primarily focused on the overall magnitude of emotional interference and have paid little attention to its temporal evolution or to the executive control mechanisms that may regulate it. In this context, transcranial direct current stimulation (tDCS) provides a valuable tool for investigating the causal contribution of prefrontal cognitive control systems to the regulation of emotional interference. The present study therefore aimed to investigate how emotional interference evolves over time and to identify the cognitive control mechanisms involved in its regulation. To address this question, we first review the prefrontal regions implicated in emotional interference and summarize evidence regarding their causal contribution and time-dependent modulation through tDCS.

### The Role of Dorsal and Ventral Prefrontal Cortex in Emotional Interference

Contemporary models propose that emotional interference emerges from the interaction between bottom-up attentional capture driven by emotionally salient stimuli and top-down control processes responsible for maintaining goal-directed behavior [9]. Within this framework, cognitive control is thought to regulate emotional distraction by maintaining task goals, prioritizing task-relevant information, and suppressing competing inputs [10,11].

Emotion–cognition interactions are thought to rely on the dynamic interplay between a ventral affective system involved in emotional appraisal and salience processing, and a dorsal executive system supporting attentional control and goal-directed behavior [9]. Consistent with this framework, neuroimaging studies have shown that successful resistance to emotional distraction is associated with increased recruitment of dorsal frontoparietal regions, particularly the dorsolateral prefrontal cortex (dlPFC), whereas emotional distraction itself is accompanied by greater engagement of ventral affective regions [12,13]. These findings suggest that the dlPFC plays a key role in maintaining task goals and reallocating attentional resources in the presence of emotionally salient distractors.

Although research on emotional interference has traditionally focused on the contribution of the dlPFC and broader frontoparietal control networks, recent evidence suggests that ventral regions may also contribute to emotion–cognition interactions. In particular, the ventromedial prefrontal cortex (vmPFC), a region classically associated with emotional appraisal and valuation processes, has recently been implicated in the regulation of emotional interference through its interactions with the dlPFC [14]. Consistent with this view, experimental work has shown that dlPFC and vmPFC stimulation can differentially influence emotional processing. For example, using an emotional evaluation task in which participants rated the valence and arousal of emotional pictures, Nejati et al. [15] reported the dissociable effects of dlPFC and vmPFC stimulation on emotional judgments. Together, these findings suggest that emotional interference may depend on coordinated interactions between dorsal executive control and ventral affective systems. However, the respective causal contributions of these regions during cognitive control tasks involving emotional interference remain poorly understood.

However, several important questions remain unresolved. First, the respective contributions of dlPFC and vmPFC were examined using global performance measures, providing limited information regarding the temporal evolution of emotional interference. Second, although the study highlighted the importance of interactions between these prefrontal regions, it did not investigate how their influence unfolds over the course of task performance. Consequently, it remains unclear whether emotional interference is regulated by stable control mechanisms or by dynamic processes that evolve over time.

The present study addresses these limitations by combining tDCS with time-resolved analyses of emotional interference. This approach allows us not only to examine the causal contribution of the dlPFC and vmPFC to executive control under emotional distraction, but also to characterize how their influence changes throughout task performance.

### Time-Dependent Effects of Prefrontal tDCS on Emotional Interference

Transcranial direct current stimulation (tDCS) has provided valuable evidence regarding the causal contribution of prefrontal regions to the regulation of emotional interference. In particular, anodal stimulation of the dlPFC has repeatedly been shown to enhance attentional control, reduce attentional biases toward threatening information, and modulate emotional reactivity [16,17,18,19,20,21]. For example, Ironside et al. [18] reported that dlPFC stimulation abolished the typical attentional bias toward threatening faces in a dot-probe task, supporting a causal role of this region in regulating the processing of emotionally salient information. More recently, interest has also extended to ventral prefrontal regions, particularly the vmPFC, which has been implicated in emotional appraisal and regulation processes [14,15,22]. Together, these findings suggest that both dorsal and ventral prefrontal systems contribute to emotion–cognition interactions.

However, an important limitation of the current literature is that stimulation effects are typically assessed using global performance measures that implicitly assume stable effects throughout task performance. This assumption may be problematic, as growing evidence suggests that the behavioral effects of tDCS are themselves time-dependent and influenced by stimulation timing [23,24,25]. For example, Friehs and Frings [26] reported stronger working-memory improvements following pre-task stimulation than during-task stimulation, suggesting that distinct neurophysiological mechanisms may underlie offline and online stimulation effects. Emotional paradigms may particularly benefit from mixed stimulation protocols, as both preparatory and state-dependent mechanisms are likely involved in emotion regulation and attentional control. Consistent with this view, stimulation initiated shortly before and maintained during emotional tasks has been shown to enhance emotion regulation and reduce attentional bias toward threat-related stimuli [17,22]. Likewise, Ohn et al. [27] observed that cognitive benefits did not emerge immediately but progressively developed during task performance.

Consequently, both emotional interference and tDCS effects appear to evolve dynamically over time. Yet, despite these converging observations, no study has directly examined how prefrontal stimulation influences the temporal evolution of emotional interference during executive control tasks. The present study was designed to address this gap by combining dlPFC and vmPFC stimulation with time-resolved analyses of emotional interference. We investigated the temporal dynamics of emotional interference under tDCS using a unified framework targeting updating and shifting processes, combined with a comparative dlPFC/vmPFC approach. Participants completed a 2-back task and a Set-switching task under neutral and negative emotional conditions while receiving 30-minute stimulation over either the dlPFC or vmPFC. The study pursued three main objectives: (i) determine how dlPFC and vmPFC stimulation modulate executive performance under emotional conditions, (ii) examine how these effects evolve over time, and (iii) assess whether stimulation timing modulates these effects.

Based on previous findings, we first expected negative emotions to impair executive control performance, particularly in cognitively demanding conditions such as non-match and switch trials, reflecting increased emotional interference [2,3,12]. Second, we hypothesized that stimulation of prefrontal regions would differentially modulate emotional interference. Specifically, dlPFC stimulation was expected to facilitate executive control of emotional distraction, resulting in reduced emotional interference relative to sham stimulation, consistent with evidence showing that dlPFC stimulation enhances executive control and reduces attentional biases toward emotional information [17,18,28,29]. In contrast, vmPFC stimulation was expected to primarily influence the processing and appraisal of emotional salience [15,22], leading to a distinct pattern of modulation. Accordingly, we predicted the dissociable effects of dlPFC and vmPFC stimulation on emotional interference. Third, because both emotional interference and tDCS-induced neuromodulation appear to evolve over time rather than remain stable throughout task performance [27,30], we hypothesized that stimulation effects would emerge progressively as stimulation exposure increased. Finally, given the limited evidence directly comparing stimulation timing protocols, exploratory analyses were conducted to examine whether the temporal profile of stimulation effects differed between online and pre-task stimulation conditions.

## Method

### Ethics Statement

This experiment received approval from *Comité de Protection des Personnes Sud-Est IV in France* (Ref #: 2023-A01670-45).

### Sample Size

Because no previous study had investigated the temporal dynamics of tDCS effects on emotion– cognition interactions across executive control processes, sample size estimation was based on the primary hypothesis of the study, namely that prefrontal stimulation would modulate emotional interference. The expected effect size was derived from the closest available study, which reported a large Emotion × Stimulation interaction during a working-memory updating task [31]. An a priori sample size calculation was performed using G*Power [32], targeting an alpha level of .05 and a statistical power of 90 %, which indicated a minimum sample size of 36 participants. To provide a margin above this estimate and account for potential inter-individual variability in responsiveness to tDCS, 48 participants were recruited from the French Air Force Academy (16 females; mean age = 24.85 years, range = 21–38). Although the sample size calculation was based on the number of participants, the primary analyses relied on a longitudinal repeated-measures design, with repeated observations collected across stimulation time points and executive-control conditions and analysed using generalized estimating equations (GEE), which account for within-subject dependence. The subsequent time-resolved analyses were motivated by our a priori hypothesis that the effects of tDCS on emotional interference evolve dynamically over time. However, because no previous study had investigated these temporal dynamics, no empirical basis was available to perform a dedicated a priori power analysis for these analyses. Accordingly, they were designed to characterize the temporal profile of stimulation effects across the course of the task.

### Emotional Stimuli

Emotions were induced trial-by-trial using 528 pictures from the International Affective Picture System (IAPS) [33]. Neutral pictures depicted everyday objects (e.g., forks), whereas negative pictures depicted aversive content (e.g., mutilations). Neutral and negative stimuli were counterbalanced across trial type (S1 Table) and presented in a randomized order across trials.

### Transcranial Direct Current Stimulation

Stimulation was delivered using a battery-driven stimulator (NeuroConn GmbH, Ilmenau, Germany). Small electrodes (3 cm^2^) were used to increase stimulation focality and reduce current spread, consistent with recent focal tDCS approaches [34,35]. An impedance threshold was calculated as a function of electrode size and was not allowed to be exceeded in order to ensure safe stimulation conditions. Electrode impedance was continuously monitored throughout stimulation and maintained below 20 kΩ. Following the international 10–20 EEG system, two active stimulation montages were employed (Fig 1). For dlPFC stimulation, the anodal electrode was positioned over F3 [17,18,22,36], whereas for vmPFC stimulation it was positioned over Fpz [15,37]. In both montages, the cathodal electrode was placed over Cz to provide a common return electrode across stimulation conditions while maintaining comparable current flow patterns [38]. During active stimulation, a constant current of 2 mA (current density = 0.67 mA/cm^2^) was delivered for 30 min, including 30-s ramp-up and ramp-down periods. In the sham condition, stimulation consisted of a 30-s ramp-up followed by a 30-s ramp-down period, with no active stimulation thereafter. Participants were blind to the stimulation conditions. No adverse effects requiring discontinuation of stimulation were reported.

**Fig 1.**
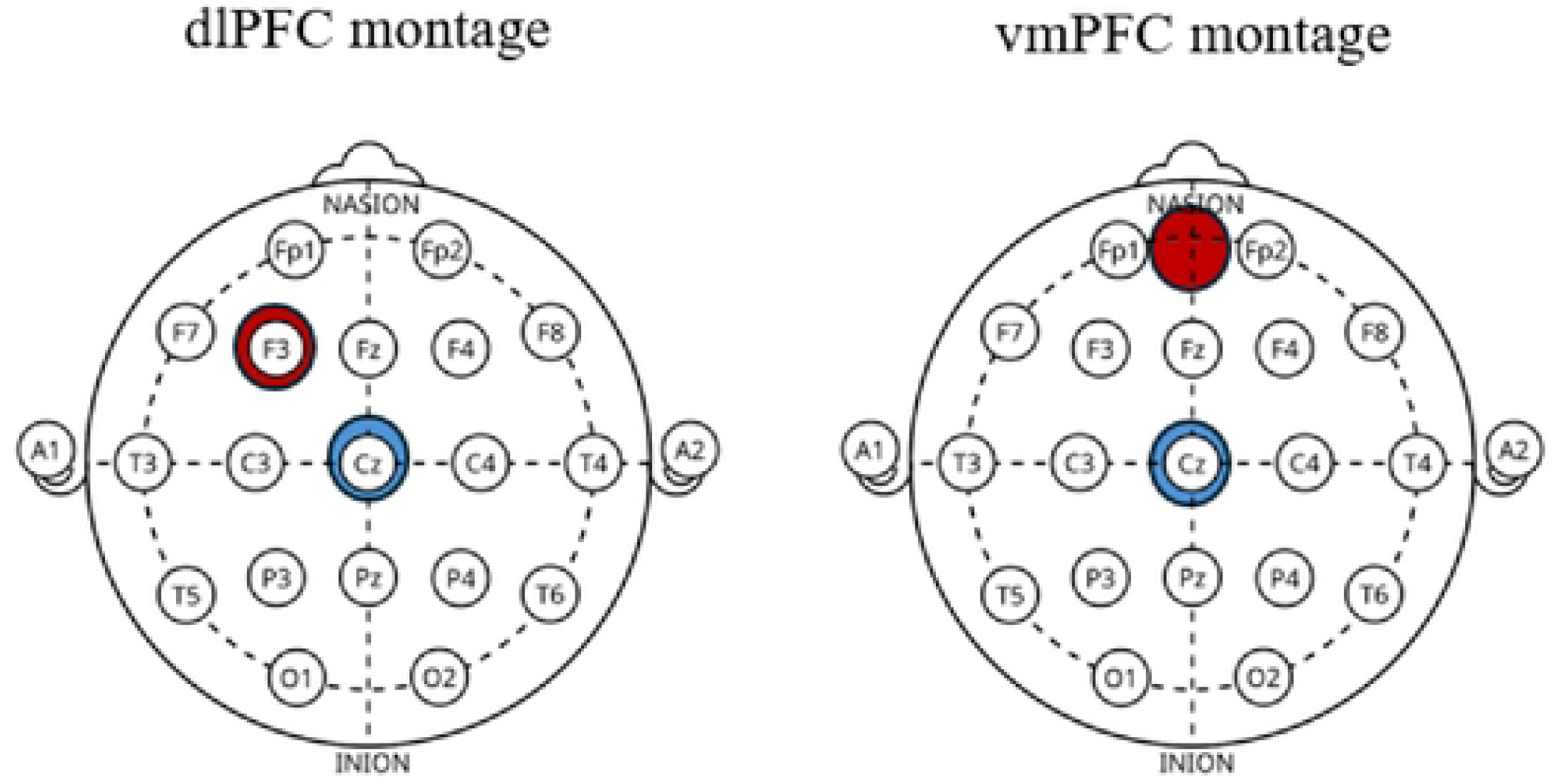
Schematic representation of the tDCS montage, with electrodes shown in red and the cathodal electrode in blue.

### Cognitive Tasks

#### Two-Back task

Participants were instructed to judge whether a digit number matched the one presented two trials earlier. The digit numbers were comprised between 1 and 9 (the ratio of odd and even numbers was counterbalanced across trial type) and were displayed on a black square, which covered about 10% of the screen. Participants were asked to press “L” key when there was a match, and “S” key when there was no match. Half of the participants received the inverse mapping. This task included four blocks of 72 trials (24 match and 48 non-match trials). Neutral pictures were displayed before half of the trials, whereas negative pictures were shown for the remaining trials. A trial was composed of a fixation cross presented for 1000 ms, followed by an emotional picture for 1000 ms, and then, a digit number was presented on the emotional pictures for 1500 ms, or until participant’s response. In order to ensure similar exposure time across participants, emotional pictures were until the end of the 1500 ms. Thus, response time variability did not imply distinct exposure time.

#### Set Switching

This task consisted of an odd and an even colored number presented one above the other. Participants were asked to determine whether the number displayed in a targeted color was odd or even, by pressing “S” key for odd numbers and “L” key for even numbers. The target color was indicated at the beginning of each block (half of the participants received the inverse mapping). Three switch-rules per block occurred (in trials 8, 17 and 24), instructing participants to process a new color. The new color became the target, while the previous one became a distractor. Participants completed 8 blocks of 30 trials (27 repeated and 3 switch trials). Neutral emotions were induced for half of the trials, and negative ones for the other half. A trial started with a fixation cross (1000 ms), followed by an emotional picture (1000 ms). Then, the digit stimuli were presented for 1000 ms, superimposed on the emotional picture. Once again, the emotional pictures were still displayed if participants respond before the end of the 1000 ms. Only the four trials preceding and following switch-rules were included in the analysis. A schematic illustration of the trial structure is presented in Fig 2.

**Fig 2.**
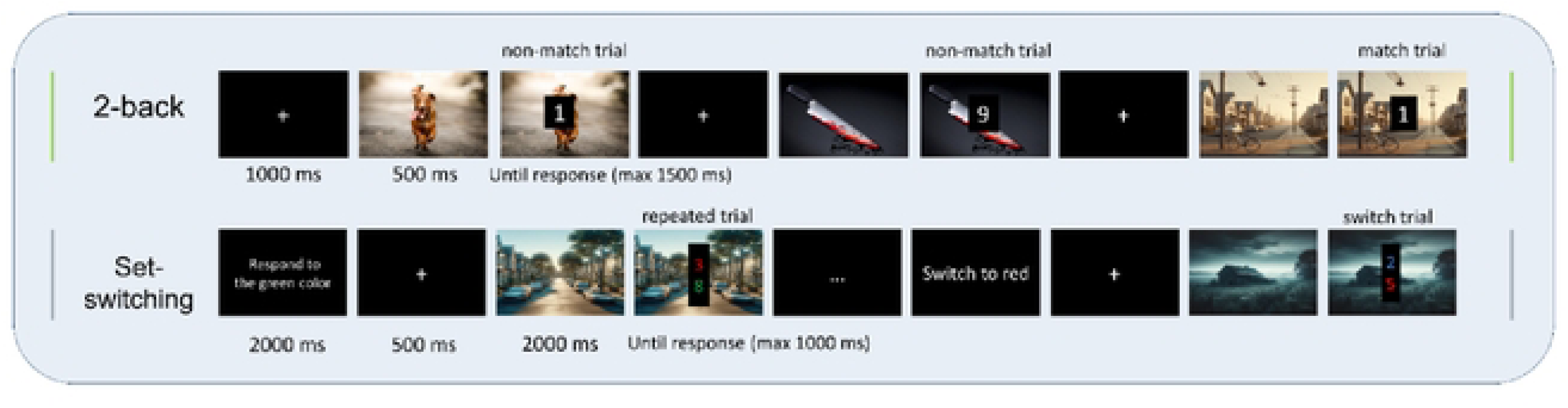
Schematic illustration of the trial sequence in the 2-back and Set-switching tasks.

### Procedure

We used a single-blind, sham-controlled, mixed design. Stimulation condition (dlPFC, vmPFC, sham) was manipulated between subjects, whereas Emotion (neutral, negative) and Trial Type were within-subject factors. All participants completed both cognitive tasks under neutral and negative emotional conditions. Within each stimulation group, participants were further assigned to one of two timing conditions: online stimulation, in which stimulation started at the onset of the first task, or pre-task stimulation, in which stimulation began 5 minutes before task onset and continued during task performance.

Participants were recruited between 27 February 2025 and 10 March 2026, and were blind to the stimulation condition. After providing written informed consent, they completed a practice phase, followed by a 30-minute stimulation period. The two cognitive tasks lasted approximately 40 minutes in total, such that stimulation covered the initial portion of task performance in both timing conditions. The task order was counterbalanced and randomized across participants.

To examine the temporal dynamics of tDCS effects, trials were divided into six consecutive time bins (Fig 3). Importantly, temporal bins were defined according to cumulative stimulation exposure rather than elapsed task time. Consequently, statistical comparisons between online and pre-task conditions were performed between bins corresponding to equivalent durations of stimulation (e.g., t5 vs. t5 = 5–10 min of cumulative stimulation), even though these bins did not necessarily correspond to the same stage of task performance. For each participant and condition, trials were ordered chronologically and grouped into six consecutive 5 minutes segments. This procedure provided a compromise between temporal resolution and statistical power while ensuring a sufficient number of observations within each bin. The resulting factor Time (t0, t5, t10, t15, t20, t25) was included in generalized estimating equation (GEE) models to capture changes in emotional interference as a function of stimulation exposure over the course of the experiment.

**Fig 3.**
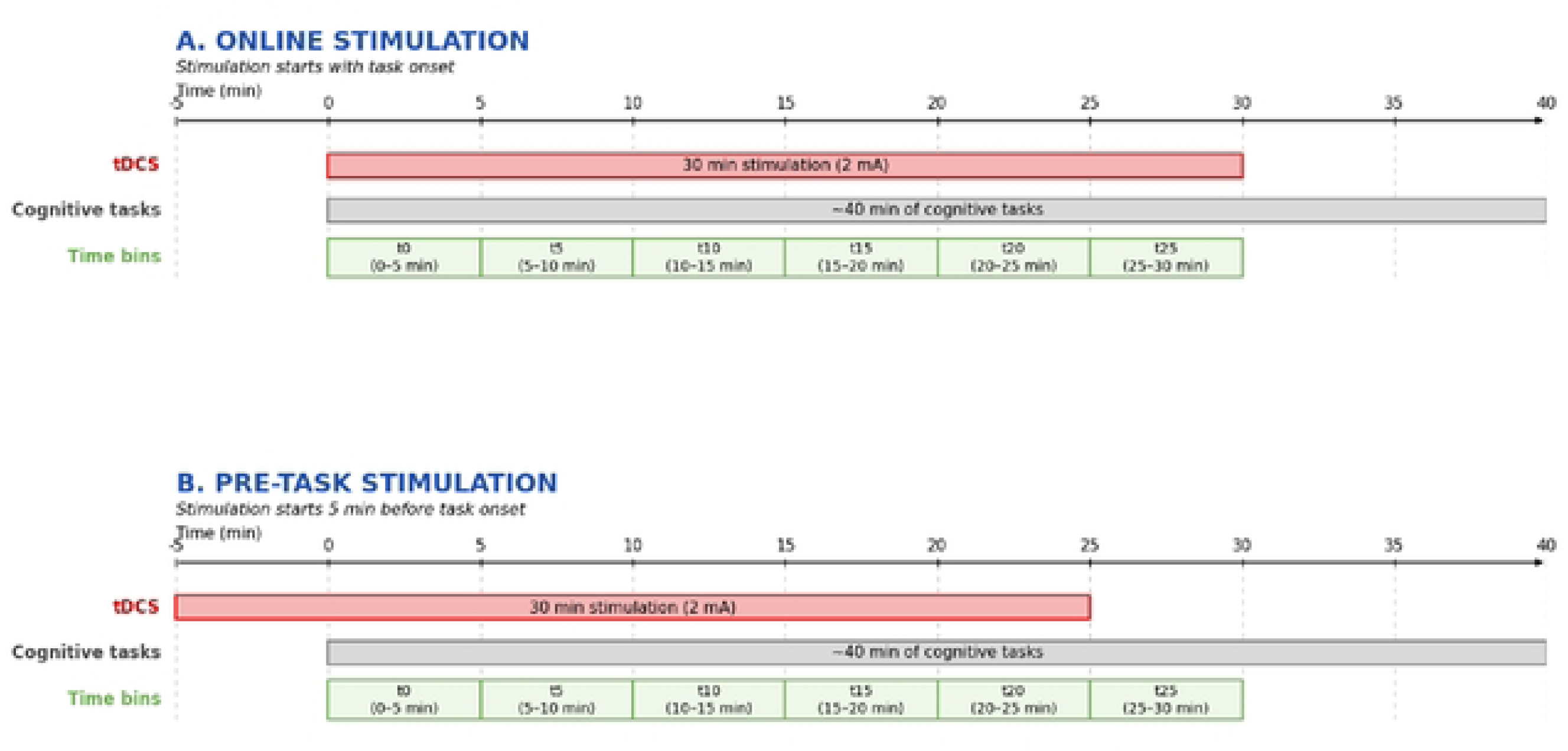
Experimental design and temporal segmentation.

### Data Analysis

Mean reaction times (RTs) and error rates were analyzed separately. To establish the presence of robust emotional interference and to validate the experimental paradigm, mixed-design repeated-measures ANOVAs were first conducted, including Emotion (neutral, negative) and Trial Type (match vs. non-match; repeated vs. switch) as within-subject factors, and Stimulation (dlPFC, vmPFC, sham) and Timing (online, pre-task) as between-subject factors. These analyses served primarily to verify that the tasks reproduced the emotion-related behavioral patterns previously reported by [39], before examining the temporal dynamics of emotional interference using time-resolved analyses. Separate ANOVAs were conducted for each task.

To further investigate how stimulation modulated emotional interference over time, an emotion-related effect score was computed for each dependent variable and trial type. Emotional effects score was calculated using the following formula:

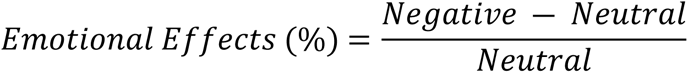

Positive values indicate greater emotional interference. This index yielded an individual-level percentage score reflecting the relative impact of negative emotions compared to neutral trials. Emotional effects score was computed separately for each trial type (match, non-match, repeated, and switch) and was used to directly quantify emotional interference. This approach allowed us to isolate performance changes specifically attributable to emotional content while reducing inter-individual variability in overall response speed. Importantly, expressing emotional effects as within-subject costs also facilitated direct comparisons across executive control tasks that differed in their baseline response times and performance characteristics.

To characterize the temporal dynamics of emotional interference, trials were ordered chronologically and divided into six consecutive 5 minute bins for each participant, yielding a within-subject factor Time (t0, t5, t10, t15, t20, t25). Because switch-trial estimates were computed from the trials immediately surrounding each rule change, fewer observations contributed to these estimates. We therefore verified observation density across groups and found no substantial imbalance in data availability between experimental conditions. Emotional costs, defined as the relative performance difference between negative and neutral trials, were computed within each time segment and used as dependent variables in subsequent analyses.

Temporal effects were examined using generalized estimating equation (GEE) models. All analyses were based on a common model structure including Trial Type, Stimulation, Timing, and Time as repeated factors, with task order included as a covariate. Because participants completed both tasks within the same stimulation session and task order was randomized, Task and Time could not be modeled independently without substantial loss of temporal information. Therefore, trial types were coded as a four-level task-specific factor (match, non-match, repeated, switch). Models included all lower- and higher-order interactions and were specified as follows:

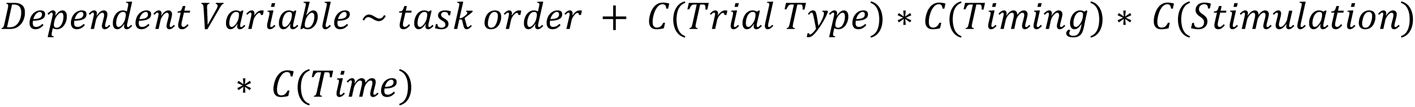

Three complementary approaches were used to characterize temporal effects: (i) baseline comparisons were conducted by treating Time as a categorical factor with t0 as the reference level, allowing each subsequent time point (t5–t25) to be directly compared to baseline: (ii) adjacent bin comparisons, in which consecutive time bins (e.g., t0 vs. t5, t5 vs. t10) were contrasted, enabling the identification of critical transitions in the evolution of emotional interference; and (iii) within-bin comparisons in which stimulation conditions (dlPFC, vmPFC, sham) were directly compared at each time point.

GEE models were fitted using an exchangeable correlation structure to account for within-subject dependencies and provide population-averaged estimates [40]. Finally, additional models included observation density as a covariate to ensure that the observed effects were not attributable to unequal numbers of observations across conditions.

## Results

For each task, latencies exceeding ± 2.5 standard deviations (SDs) were excluded, representing the removal of 2 % for the 2-Back task and 1 % for Set Switching task. Only correct trials were included in the analysis of RTs. For a summary of the results, see Table 1.

**Table 1.**
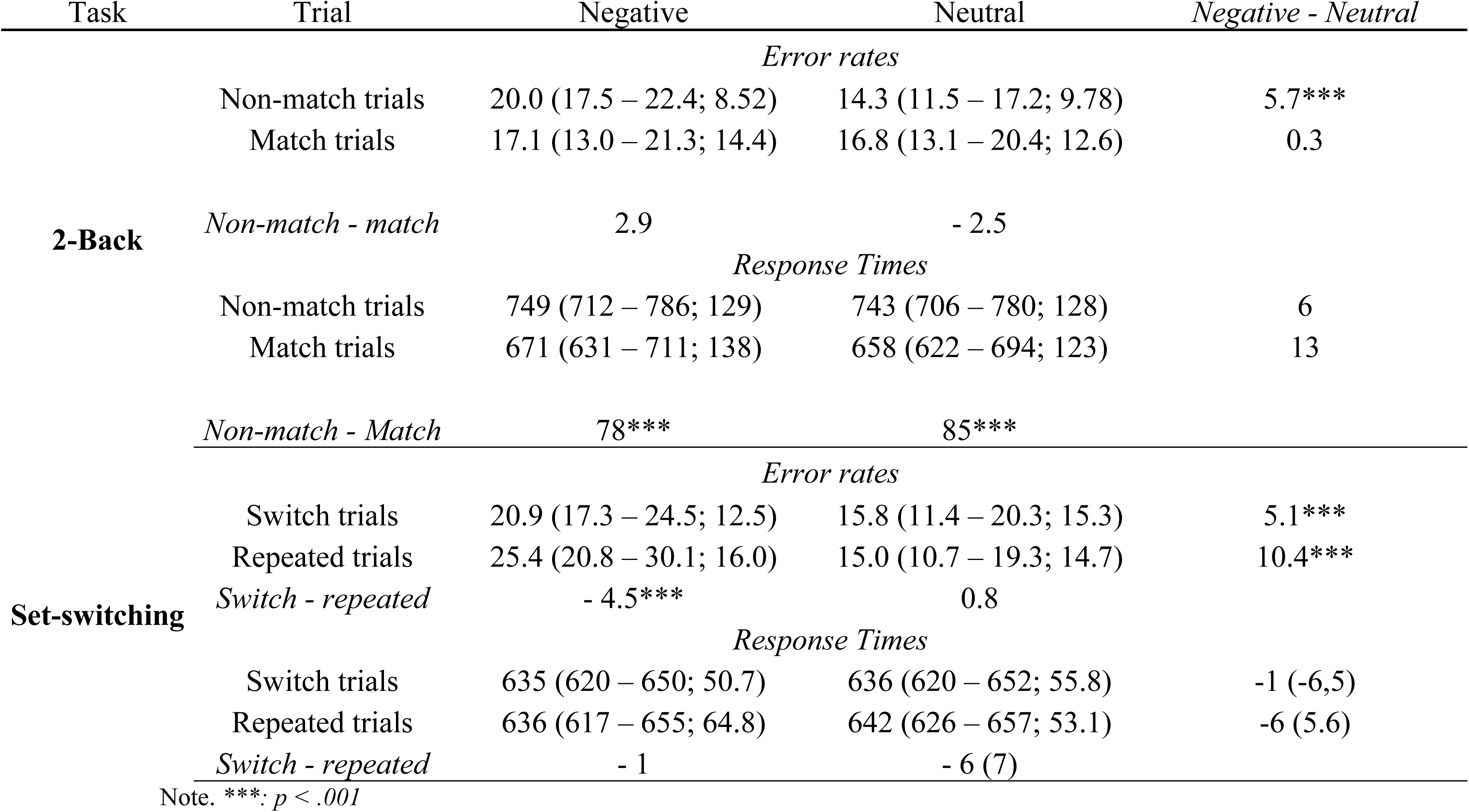
Mean error rates and response times for 2-back and Set-switching tasks in negative and neutral emotions.

### Emotional Interference Across Cognitive Control Task

#### Two-back task

RTs analyses revealed a main effect of Trial Type under neutral emotions, *F*(1, 47) = 49.0, *p* < .001, *MSe* = .08, *ƞ^2^_p_* = .55. Participants are slower to respond to non-match trials (738 ms) compared to match trials (650 ms). A similar effect also came out significant on the whole sample, *F*(1, 47) = 53.25, *p* < .001, *MSe* = .32, *ƞ^2^_p_* = .53, with increased RTs in non-match trials (743 ms) compared to match trials (658 ms). Interestingly, the analyses revealed a main effect of Emotion, *F*(1, 47) = 5.68, *p* = .021, *MSe* = .01, *ƞ^2^_p_* = .11, reflecting slower RTs under negative emotions (710 ms) compared to neutral emotions (700 ms). No main effect of Stimulation or interaction involving stimulation sites was observed. However, a main effect of Timing indicated faster responses to the online condition (664 ms) compared to the pre-task condition (732 ms) *F*(1, 47) = 5.37, *p* = .025, *MSe* = .30, *ƞ^2^_p_* = .11.

Error rate analyses revealed a main effect of Emotion, qualified by a significant Emotion × Trial Type interaction, *F*(1, 47) = 14.64, *p* < .001, *MSe* = 330.97, *ƞ^2^_p_* = .24. Post hoc comparison (Holm-Bonferroni-corrected) showed that negative emotions selectively increased error rates in non-match trials (20%) compared to neutral trials (14%), *t*(47) = 8.31, *p* < .001 whereas no reliable difference was observed for match trials. A main effect of timing was also observed, *F*(1, 47) = 5.95, *p* = .019, *MSe* = 2056, *ƞ^2^_p_* = .12, with lower error rates in the online condition (13%) compared to the pre-task condition (20%). These results indicate that emotional interference primarily affects high-demand updating processes.

#### Set-switching task

No main effect of Trial Type was observed under neutral emotion, for either response times or error rates. However, analyses of error rates across the full sample revealed a significant Emotion × Trial Type interaction, *F*(1, 47) = 6.37 *p* = .015, *MSe* = 346.00, *ƞ^2^_p_* = .12. Post hoc comparisons indicated that the difference between repeated and switch trials emerged only under negative emotions, *t*(47) = 2.43, *p* = .038, whereas no reliable difference was observed under neutral conditions, *t*(47) = −1.05, *p* = .301. This pattern indicates that task-related differences became behaviorally apparent, specifically in the presence of emotional interference.

Crucially, no global effects of tDCS were observed in these ANOVA analyses. Therefore, we next examined whether stimulation effects emerged over time using time-resolved GEE models.

### The Effects of tDCS on Emotional Interference Across Time

To investigate whether tDCS modulated emotional interference as a function of stimulation exposure, generalized estimating equation (GEE) models were applied to the emotional cost indices derived from both RTs and error rates. Because participants completed both the 2-back and Set-switching tasks within a single experimental session, data from both tasks were analyzed within a common modeling framework, with the task order included as a covariate.

The complete model outputs are provided in Table A2.

### Baseline

In the sham condition, emotional interference on RT decreased over time, reflecting a gradual attenuation of emotional effects on match trials during task performance. Relative to baseline (t0), emotional costs were significantly reduced at t5 (β = −.14, SE = .04, 95 % CI [−.22, −.06], p = .001), t10 (β = −.17, SE = .07, 95% CI [−.31−.03], p = .015), and t15 (β = −.19, SE = .07, 95% CI [−.33, −.05], p = .008). No further changes were observed at later time points (i.e., t20 and t25). The temporal evolution of RT emotional effects on match trials is illustrated in Fig 4.

**Fig 4.**
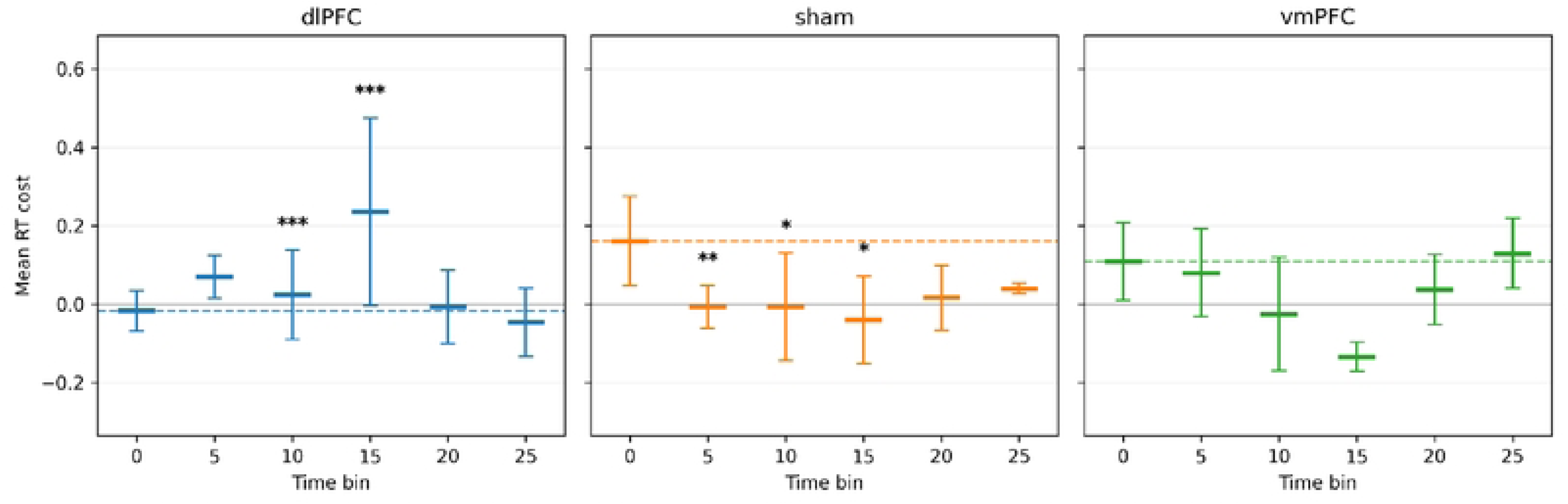
Time-resolved RT emotional effects in match trials during online stimulation. Horizontal segments represent mean RT emotional costs for each 5-min time bin, with 95% confidence intervals. Dashed lines indicate baseline (t0) emotional costs. Asterisks denote significant effects identified by GEE analyses and do not necessarily reflect descriptive differences visible in the plotted means.

In contrast, active stimulation modified this temporal pattern. Significant stimulation-related effects emerged around t10, indicating that the temporal evolution of emotional interference differed from that observed in the sham condition. These effects varied as a function of both stimulation sites and trial type.

The largest effects were observed following dlPFC stimulation, with significant modulation of emotional costs around t10 for both match (β = .61, SE = .13, 95% CI [.36, .86], p < .001) and non-match trials (β = .23, SE = .12, 95% CI [−.01, .47], p = .044), particularly in the online condition. Follow-up baseline comparisons further showed that the increase in emotional costs observed for match trials under dlPFC stimulation remained significant at t15 relative to baseline (β = .23, SE = .02, 95% CI [.19, .27], p < .001). In comparison, vmPFC stimulation produced more selective effects, primarily affecting switch trials (β = −.43, SE = .06, 95% CI [−.55, −.31], p < .001).

Overall, these findings indicate that the effects of tDCS on emotional interference are time-dependent, emerging progressively and reaching their greatest magnitude approximately 10 minutes after stimulation onset.

### Adjacent bin

Analyses of consecutive time bins revealed that the most pronounced stimulation-related changes in emotional interference occurred between 5 and 15 minutes of cumulative stimulation exposure. dlPFC stimulation modulated emotional effects early, with significant changes emerging between t5 and t10 for match trials (β = −.48, SE = .18, 95% CI [−.83, −.13], p = .007) and persisting between t10 and t15 (β = .59, SE = .24, 95% CI [.12, 1.06], p = .012). In contrast, vmPFC effects emerged later and were more selective, primarily affecting switch (β = −.08, SE = .03, 95% CI [−.14, −.02], p = .018) and non-match trials (β = −.10, SE = .05, 95% CI [−.20, −.00], p = .037) between t10 and t15. Importantly, these effects were transient and largely disappeared after 20 minutes, with only a small residual modulation observed for match trials under dlPFC stimulation between t20 and t25 (β = −.11, SE = .05, 95% CI [−.21, −.01], p = .039).

Together, these findings suggest distinct temporal profiles for dlPFC and vmPFC stimulation, with earlier and more sustained effects following dlPFC stimulation and later, more selective effects following vmPFC stimulation.

### Within-bin comparison

Direct comparisons within each time bin confirmed the transient nature of tDCS effects. Significant differences between active stimulation and sham were observed at specific time points, indicating that stimulation effects were not sustained across task but instead occurred intermittently.

These effects varied as a function of the stimulation site and trial type. Differences were observed during the earliest stimulation interval (t0) for switch trials following both dlPFC (β = −.6, SE = .30, 95% CI [−1.19, −.01], p = .042) and vmPFC stimulation (β = −.87, SE = .42, 95% CI [−1.69, −.05], p = .040) (Fig 5). Additional effects emerged at t5, where vmPFC stimulation was associated with increased emotional interference (β = .50, SE = .18, 95% CI [.15, .85], p = .005), and at later stages of the experiment, including modulation of match trials following dlPFC stimulation at t20 (β = .51, SE = .18, 95% CI [.16, .86], p = .004). Overall, these findings indicate that tDCS effects were time-dependent and non-uniform, emerging at discrete moments rather than continuously throughout task performance. This pattern further supports the view that stimulation effects evolve dynamically over the course of the experiment. The descriptive results of RT emotional effects according to stimulation condition are provided in S2 Table.

**Fig 5.**
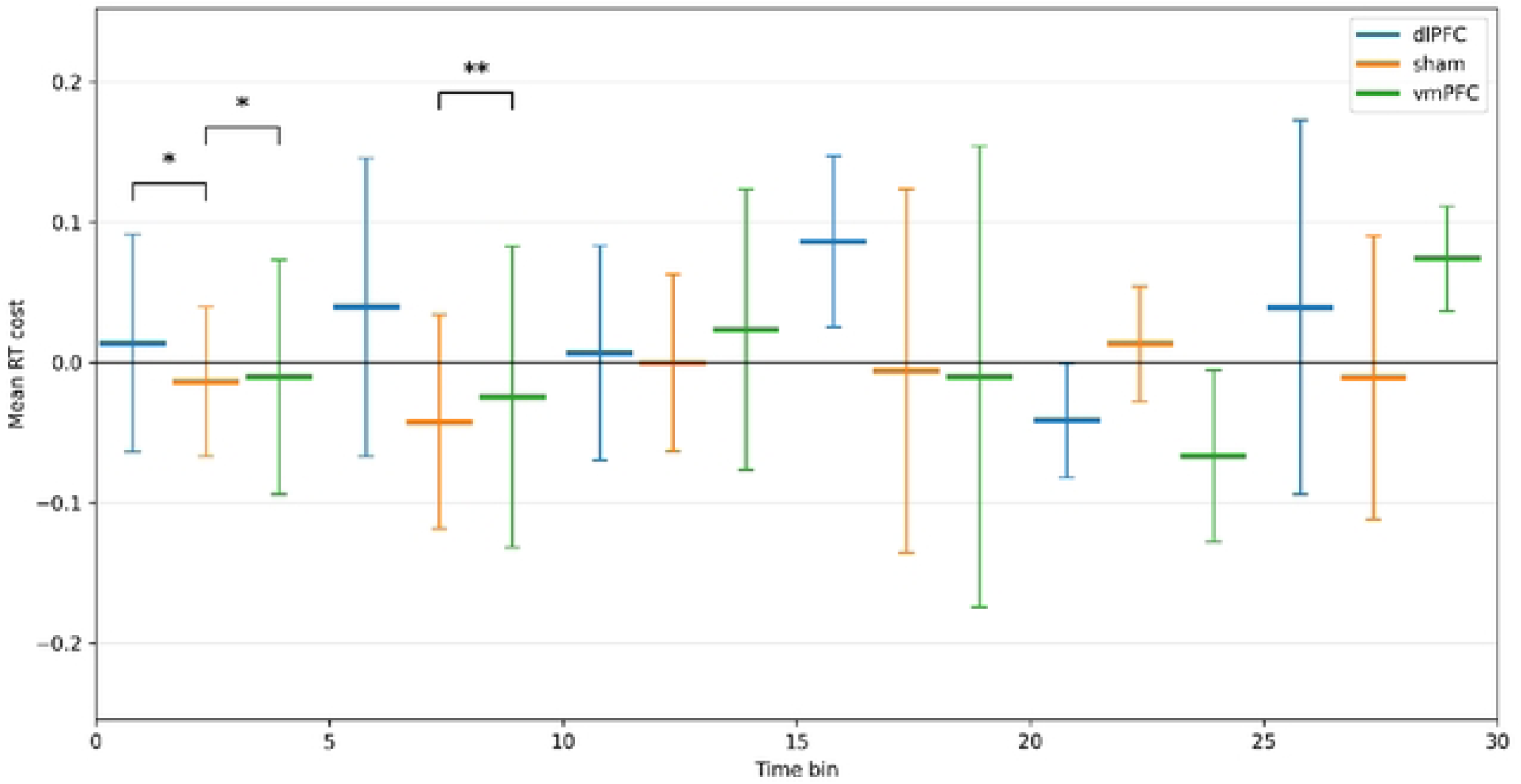
RT emotional costs across stimulation conditions for online switch trials. Asterisks indicate significant between-stimulation effects identified by GEE analyses and do not necessarily reflect differences apparent in the descriptive means displayed in the figure.

### Control analyses

Control analyses indicated that the number of observations per cell did not account for the results. The density covariate was not significantly associated with accuracy costs and showed only a marginal effect on RT costs. Importantly, its interaction with time was not significant, including t15, and the overall pattern of results remained unchanged when controlling for data density. These findings suggest that the observed effects are unlikely to be driven by unequal data distribution across time points.

## Discussion

The present study investigated how dlPFC and vmPFC stimulation modulate emotional interference during executive control performance and, critically, how these effects evolve over time. Three main findings emerged. First, the results replicated previous evidence showing that negative emotions impair executive control performance, particularly under high cognitive demands [39]. Second, no global effects of stimulation were detected using conventional analyses. Third, and most importantly, time-resolved analyses revealed that both emotional interference and stimulation effects evolved dynamically throughout task performance, with the most consistent stimulation effects emerging after approximately 10–15 minutes of cumulative stimulation exposure.

Consistent with previous findings, participants exhibited slower responses for non-match than match trials in the 2-back task, reflecting the greater demands associated with updating processes [41,42]. Likewise, negative emotions selectively impaired updating performance, as reflected by increased error rates in non-match trials. In the Set-switching task, emotional interference was associated with increased errors across both repeated and switch trials, suggesting a broader disruption of cognitive control processes. Although the classical switching cost was not observed in the present study, the overall pattern of emotional interference replicated our previous findings [39]. Contrary to our initial predictions, active stimulation did not produce global improvements in executive performance nor a generalized reduction in emotional interference. However, analyses that explicitly considered the temporal evolution of performance revealed a different pattern. dlPFC stimulation produced the most consistent modulation of emotional interference, with effects emerging approximately 10–15 minutes after stimulation onset and varying according to trial type. In contrast, vmPFC stimulation produced fewer and more selective effects, primarily affecting error rates. Importantly, these stimulation effects were transient rather than sustained, varying across temporal bins and stimulation conditions. These findings suggest that the influence of prefrontal stimulation on emotion– cognition interactions may be better understood as a dynamic process unfolding over time rather than as a stable modulation of performance. More broadly, they highlight the importance of considering the temporal evolution of both emotional interference and neuromodulatory effects when investigating the neural mechanisms underlying executive control under emotional conditions.

### How Emotions Influence 2-back and Set-switching Performance

Our findings indicate that negative emotions differentially affect updating and shifting performance, consistent with our previous work [39]. Emotional interference was particularly pronounced in updating, with specific impairment in non-match trials, whereas shifting performance was more broadly affected.

In the 2-back task, participants responded more slowly to non-match than match trials under neutral conditions, confirming the greater cognitive demands associated with updating processes [41,42]. More importantly, negative emotions selectively increased errors in non-match trials. Because these trials require continuous monitoring, comparison of incoming information with maintained representations, and suppression of no-longer-relevant information, they place greater demands on executive resources. The selective emotional impairment observed in these trials therefore supports the view that emotional stimuli compete with task-relevant processing for limited cognitive control resources, resulting in greater interference under conditions of high executive demand.

A different pattern emerged in the Set-switching task. Although the classical switching cost was not observed, negative emotions increased error rates across both repeated and switch trials. The absence of a switching cost may reflect the fixed preparation interval introduced in the present design, which may have encouraged a more proactive mode of control [43] and reduced behavioral differences between trial types. Nevertheless, emotional interference remained robust. Importantly, unlike our previous study, emotional effects were observed on accuracy but not on response times. Within a proactive-control framework, emotional stimuli may have disrupted the maintenance of task goals and task representations rather than the switching process itself [44].

These findings suggest that emotional interference does not affect all executive functions in the same manner. Rather, its impact appears to depend on the specific cognitive operations required by the task, with the strongest effects emerging when executive demands place substantial pressure on limited cognitive control resources.

### The Effects of tDCS on Emotional Interference Over Time

A key finding of the present study is that emotional interference and stimulation effects evolved dynamically over time. While conventional analyses failed to reveal robust global effects of stimulation, time-resolved analyses showed that both the magnitude of emotional interference and its modulation by tDCS varied across the course of task performance.

In the sham condition, emotional interference progressively decreased over time, particularly for match trials. Although this pattern may partly reflect habituation to repeated emotional stimuli, a pure habituation-based account appears insufficient. Participants were exposed to the same emotional material throughout the experiment, yet the temporal reduction was not observed uniformly across trial types. Instead, emotional interference decreased primarily in conditions associated with lower cognitive demands. This finding suggests that the reduction of emotional interference may reflect the progressive recruitment of adaptive control mechanisms rather than simple emotional desensitization. One possible explanation is that lower-demand trials leave greater executive resources available for the regulation of emotional distraction. This interpretation is consistent with evidence indicating that emotion regulation depends on the availability of cognitive control resources and becomes less efficient under high cognitive load [45,46,47]. Under this view, the temporal reduction of emotional interference may reflect a gradual optimization of control strategies that emerge as participants adapt to both the task and the emotional context.

Crucially, active stimulation modified this temporal trajectory. The most consistent effects emerged approximately 10–15 minutes after stimulation onset, suggesting that tDCS did not exert an immediate influence on emotional interference but progressively altered the way cognitive control mechanisms were deployed over time. This delayed emergence is consistent with evidence showing that behavioral effects of tDCS often become detectable gradually, as stimulation-induced changes in cortical excitability accumulate [27].

Interestingly, stimulation effects did not primarily manifest as global improvements in performance. Instead, they altered the temporal evolution of emotional interference. This distinction is important because it suggests that tDCS may influence the way emotional interference unfolds during task performance rather than executive performance per se. In other words, stimulation appears to modify how individuals adapt to emotional distraction over time rather than simply enhancing cognitive efficiency. Although dlPFC stimulation produced the most consistent effects, particularly during updating-related conditions, these effects were transient and dependent on task demands. The greater sensitivity of updating-related conditions to dlPFC stimulation is broadly consistent with previous work linking updating processes to prefrontal control mechanisms [12,48,49]. By contrast, vmPFC stimulation was associated with fewer and more selective effects. However, given the limited and context-dependent nature of these findings, the present results should not be interpreted as evidence for distinct functional contributions of the dlPFC and vmPFC. Rather, they suggest that the influence of prefrontal stimulation on emotional interference may vary according to task demands and the temporal window during which performance is assessed. Overall, this pattern is consistent with the view that emotional interference emerges from dynamic interactions between executive control and affective systems, whose relative contribution may vary across contexts and stages of task performance [50,51].

Taken together, the present findings suggest that emotional interference is not a static phenomenon and that its modulation by tDCS may depend on when cognitive performance is measured. More broadly, they indicate that time-resolved approaches can reveal effects that remain undetected when emotional interference and stimulation effects are assessed using global performance measures alone.

### The Influence of the Timing of Stimulation

The present findings further suggest that stimulation timing may influence the modulation of emotional interference. First, stimulation-related effects were observed more frequently in the online than in the pre-task condition. This pattern is consistent with state-dependent accounts of non-invasive brain stimulation, which propose that tDCS preferentially modulates neural processes that are actively engaged at the time of stimulation [52,53]. Because emotional interference and its regulation emerge during task performance, online stimulation may have interacted more directly with the prefrontal mechanisms involved in controlling emotional distraction. This interpretation is further supported by evidence showing that the prefrontal cortex plays a central role in the top-down regulation of emotional information and that tDCS can selectively influence emotion-related cognitive processing [22,54]. In contrast, stimulation delivered before task engagement may have induced more general changes in cortical excitability without specifically targeting the emotional and control processes recruited during task execution.

Second, stimulation effects did not emerge immediately but became detectable only after approximately 10 min of stimulation. This delayed onset is consistent with evidence indicating that behavioral effects of tDCS often require a sufficient accumulation of stimulation-induced changes in cortical excitability before influencing performance [27]. Importantly, this temporal profile closely mirrors the progressive evolution of emotional interference observed throughout the task, suggesting that stimulation interacted with adaptive control processes as they unfolded rather than exerting a direct influence from task onset.

Finally, stimulation effects appeared to be transient rather than sustained throughout the entire task. Rather than producing a stable enhancement of cognitive control, tDCS seemed to modulate specific phases of the adaptation process. This transient profile further supports the view that stimulation effects depend on the current functional state of the cognitive system and may be most effective when ongoing emotional and executive control processes are actively engaged. Together, these findings suggest that the impact of tDCS on emotional interference depends not only on where stimulation is applied, but also on when stimulation occurs relative to the dynamic recruitment of affective and control mechanisms. Our findings may help explain why previous tDCS studies have reported inconsistent effects on emotional and cognitive performance when stimulation timing was not explicitly considered.

## Conclusions

The present study demonstrates that emotional influences on cognitive control are not static but evolve dynamically throughout task performance. While negative emotions differentially impaired updating and shifting processes, tDCS effects were not observable at a global level and only emerged when the temporal evolution of emotional interference was taken into account. Specifically, stimulation effects appeared selectively, varied as a function of cognitive demands, and depended on both the stimulation site and timing. Although dlPFC stimulation produced the most consistent effects, particularly during updating-related conditions, the overall pattern suggests that emotional interference and its regulation cannot be fully understood using static measures of performance alone. More broadly, the present findings highlight the importance of considering the temporal dynamics of both cognitive and affective processes, suggesting that these dynamics may constitute a critical dimension for understanding how the brain regulates emotional influences on cognition.

Several limitations should be acknowledged. First, although the time-resolved analyses suggest that tDCS effects on emotional interference emerged dynamically over the course of the session, these temporal effects cannot be attributed exclusively to stimulation. They may also partly reflect habituation to emotional stimuli, fatigue, task learning, or the progressive implementation of adaptive strategies. Future studies should, therefore, include additional control conditions or designs allowing these temporal factors to be disentangled from stimulation-related effects. Second, stimulation conditions were manipulated between participants. Given the substantial interindividual variability in tDCS responsiveness, a within-subject or cross-over design would provide a stronger test of stimulation effects by reducing between-subject variability. Third, no clear stimulation effect was observed in switch trials. This may be due to the relatively transient nature of the switching process in the present task, or to the limited number of switch trials available for time-resolved analyses. Future work could use tasks that place more sustained demands on shifting or task-set reconfiguration, which may be better suited to reveal dlPFC contributions. Fourth, although the Fpz-Cz montage was chosen based on previous work targeting vmPFC-related processes, no electric-field modeling was performed. Therefore, the anatomical specificity of the stimulation remains uncertain and interpretations regarding vmPFC involvement should be made with caution. Finally, although GEE models provided stable population-level estimates for the present complex repeated-measure structure, linear mixed-effects models would have offered a more flexible way to model participant-level variability. However, such models would need to be sufficiently stable and convergent to support reliable interpretation.

## Acknowledgement

We gratefully acknowledge the participation and contribution of all individuals who took part in this study. The authors also would like to thank Dr. Laurence Casini and Dr. Sébastien Scannella for their helpful comments and advice on the tDCS methodology used in this study.

## Data availability statement

The data and code supporting the findings of this study are openly available in Open Science Framework (OSF): https://doi.org/10.17605/OSF.IO/6BCYP

## Supporting Information

**S1 Table.** Mean emotional valence and arousal (confidence interval and standard deviation) for each trial type.

**S2 Table.** Mean emotional effects (standard deviation) on response times across Trial Type, Stimulation, Time bins, and Timing condition

## Author Contributions

Tristan Feutren: conceptualization, investigation, writing – original draft, methodology, visualization, writing – review and editing, formal analysis, data curation, and software. Valentin Braud: writing – review and editing, software, data curation. Ludovic Fabre: Conceptualization, writing – review and editing, supervision, project administration.

